# Genome-wide correlation analysis reveals *Rorc* as potential amplitude regulator of circadian transcriptome output

**DOI:** 10.1101/2020.06.19.161307

**Authors:** Evan S. Littleton, Shihoko Kojima

## Abstract

Cell-autonomous circadian system, consisting of core clock genes, generates near 24-hour rhythms and regulates the downstream rhythmic gene expression. While it has become clear that the percentage of rhythmic genes varies among mouse tissues, it remains unclear how this variation can be generated, particularly when the clock machinery is nearly identical in all tissues. In this study, we sought to characterize circadian transcriptome datasets that are publicly available and identify the critical component(s) involved in creating this variation. We found that the relative amplitude of 13 genes and the average level of 197 genes correlated with the percentage of cycling genes. Of those, the correlation of *Rorc* in both relative amplitude and the average level was one of the strongest. In addition, the level of *Per2AS*, a novel non-coding transcript that is expressed at the *Period 2* locus, was also linearly correlated, although with a much lesser degree compared to *Rorc*. Overall, our study provides insight into how the variation in the percentage of clock-controlled genes can be generated in mouse tissues and suggests that *Rorc* and potentially *Per2AS* are involved in regulating the amplitude of circadian transcriptome output.

## Introduction

Circadian clocks regulate the daily fluctuations of biochemical, physiological, and behavioral rhythms ^1^. In mammals, signals originating in the suprachiasmatic nucleus (SCN) of the hypothalamus synchronize independent oscillators in other peripheral tissues, such as the brain and even in fibroblasts ^2,3^. The molecular circadian clock within each cell is comprised of interlocking transcriptional-translational feedback loops, whose coordinated action is essential to generating cell-autonomous circadian oscillation ^4^.

At its core mechanism, the BMAL1 (official gene name: *Arntl*) and CLOCK (or its paralogue NPAS2) form a heterodimer and activate the transcription of *Period (Per) 1-3* and *Cryptochrome* (*Cry)1-2*, whose promoters contain target DNA regulatory elements, called E-boxes. As the level of PER and CRY proteins increases, they form a heterodimer and translocate back to the nucleus to repress their own transcription. As repression of *Per* and *Cry* transcription progresses, the level of the PER/CRY protein decreases, thereby allowing ARNTL and CLOCK to begin a new cycle of transcription. As an auxiliary loop, ARNTL and CLOCK also activate the expression of *Rev-erba/b* (official gene name: *Nr1d1* and *Nr1d2*), and *Rora-c*, all of which are nuclear receptors. REV-ERB and ROR proteins, in turn, repress or activate the target mRNA expression including *Arntl, Clock*, and *Npas2*, respectively, by recognizing DNA elements termed REV-ERB/ROR binding motifs (ROREs) in their promoters. As an additional loop, ARNTL/CLOCK activates the expression of *Dbp*, which activates the transcription of target mRNAs that possess a DNA element, called a D-box, while REV/ROR proteins regulate the expression of *Nfil3*, which represses D-box containing genes. Targets include *Rev-erbs, Rors*, and *Pers* ^4^.

Cell-autonomous circadian clocks also drive thousands of rhythmic output genes (i.e., clock-controlled genes) that, ultimately, produce daily rhythms of many types of physiology and behavior ^5-10^. Interestingly, the number of cycling genes is vastly different among mouse tissues. In some tissues, more than ten percent of the entire transcriptome is rhythmic, while only a few percent are rhythmic in other tissues ^10-12^. Nevertheless, it remains unclear how the core clock machinery drives different numbers of clock-controlled genes, even though the core clock mechanism is nearly identical in each tissue.

To gain mechanistic insights into how some tissues produce more cycling genes than others, we characterized the circadian transcriptome data from various mouse tissues and attempted to identify a parameter(s) that correlates with the percentage of clock-controlled genes. We found that the differences in the percentage of cycling genes are not due to the strength of transcriptome. Interestingly, however, the relative amplitude of 13 genes as well as the average level of 197 genes correlated with the percentage of cycling genes. Of our particular interest was *Rorc*, whose correlation in both the relative amplitude and the average level was one of the strongest. We also found that the level of *Per2AS*, a novel non-coding transcript that is expressed at the *Period 2* locus, also showed a correlation. Based on these data, we propose that *Rorc* is involved in regulating the amplitude of circadian transcriptome output, although the effect of *Rorc* is most likely independent of its activity as a transcriptional activator, as the percentage of cycling genes with RORC binding motif in their promoter was consistent across all the tissues.

## Methods

### Microarray Data Processing

Microarray data were downloaded through NCBI GEO from series GSE54650 ^10^. Data was originated from 12 different tissues, with 24 time points from each tissue in 2-hour intervals over the course of 48 hours. Extracted data was normalized by Robust Multichip Average (RMA) normalization ^16^ and annotated by the Affymetrix Transcriptome Analysis Software package (http://www.affymetrix.com/support/technical/byproduct.affx?product=tac). Unannotated probesets, as well as those that had values lower than the average of all negative probesets across all timepoints in the respective tissue, were eliminated from the downstream analysis. For multiple probesets annotated to the same gene, the probeset with the highest average value was selected.

### RNA-seq Data Processing

RNA-seq data were downloaded as fastq files through the NCBI database from SRA ID SRP036186 ^10^. Data contained information from 12 different tissues, with 8 time points from each tissue in 6-hour intervals over the course of 48 hours. Reads were mapped to the Ensembl mouse genome release 95 using STAR 2.7.0a ^17^ with outFilterScoreMinOverLRead = 0.3 and outFilterMatchNMinOverLRead = 0.3 options. We added the option ‘Condensegenes’ to select the most abundant isoform as the representative of a gene, as well as the option ‘count exons’ to measure only mRNA. The quantification of expression level was performed by HOMER ^18^ using the transcripts per million (TPM) option. Any transcript with an average TPM < 0.5 across all timepoints were eliminated from the downstream rhythmicity analysis. We also used TPM to normalize the expression levels of each transcript. We eliminated white adipose data from the downstream analysis because no transcripts were rhythmic with our statistical threshold (BH. Q-value <0.05), even though more than 13,500 transcripts were detected after applying the filter of TPM > 0.5. The expression of *Per2AS* was measured with the “strand –” option in HOMER. We did not apply the filter (TPM > 0.5 to call ‘expressed’) in quantifying the level of *Per2AS*, because non-coding transcripts generally have low expression levels ^19-21^.

### Rhythmicity Analysis

We used MetaCycle ^22^ to determine the rhythmicity of each gene. MetaCycle integrates three different algorithms ARSER, JTK_CYCLE, and Lomb-Scargle and calculates the p-value, Benjamini-Hochberg q-value (BH.Q value), period, phase, baseline value, amplitude (AMP), and relative amplitude (rAMP), which is the ratio between amplitude and baseline expression level. We defined the expression rhythmic when meta2d BH.Q < 0.05.

### Correlation Analysis

Pearson and Spearman correlation tests were performed in R to determine the linear and non-linear correlation between the percentage of cycling transcripts in each tissue and the rhythmicity (using BH.Q value), phase, and relative amplitude of the 15 clock genes, as well as *Per2AS*, calculated by the MetaCycle package in R. A significant correlation was defined as a p-value < 0.05. For the transcriptome-wide correlation analysis, we used the rcorr function from the Hmisc and tidyverse packages in R to perform Pearson or Spearman correlation tests and used the average gene expression of transcripts expressed in all 12 (microarray) or 11 tissues, excluding white adipose tissue (RNA-seq)^23,24^. Fisher Z-scores were calculated from the Rho or R^2^ with Fisher transformation. GO enrichment analysis of the significantly correlated genes was performed using the Gene Ontology Resource ^25,26^.

### Promoter Motif Analysis

We first retrieved the promoter sequences (−1000 bp to +100 bp with respect to the transcription start site: TSS) of all the cycling transcripts from the UCSC Genome Browser, and performed a motif search using Find Individual Motif Occurrence (FIMO) with a p-value = 1×10^− 4^ as the threshold ^27^. Input motif matrices were downloaded from JASPER (RORA: MA0071.1, RORA(var.2): MA0072.1, RORB: MA1150.1, RORC: MA1151.1, NR1D1: MA1531.1, NR1D2: MA1532.1, ANRTL: MA0603.1, CLOCK: MA0819.1, NPAS2: MA0626.1, NFIL3: MA0025.1, and MA0025.2, DBP: MA0639.1) ^28^.

## Results

### Characteristics of circadian transcriptomic output in various mouse tissues

To gain insight into what determines the number of clock-controlled genes in each tissue, we first retrieved existing circadian transcriptome datasets from various mouse tissues ^10^. We found this particular dataset best-suited to our study, because it covered the highest number of tissues (12 total) and provided the highest time resolution (2hr intervals), compared to other studies ^5,7-9^.

Our in-house analysis was able to replicate the previous findings, in which the percentage of cycling genes was highest in liver, followed by kidney, lung, brown adipose, and heart, and lowest in brainstem (Fig. 1A). The ranks are slightly different from the original study ^10^, which is most likely due to the differences in the analytical methods and statistical criteria used in our study (see Materials and Methods). Distribution of Benjamini-Hochberg q-values from the rhythmicity analysis was also widest in liver, followed by kidney, lung, brown adipose, and heart, and was particularly narrow in white adipose and brainstem (Fig. 1B). These data indicate that in liver, the expression of many transcripts fluctuates even if it does not reach a statistically significant level for circadian rhythmicity, while in white adipose and brainstem, the expression of the majority of the transcripts do not fluctuate at all (Fig. 1B).

**Figure 1.**
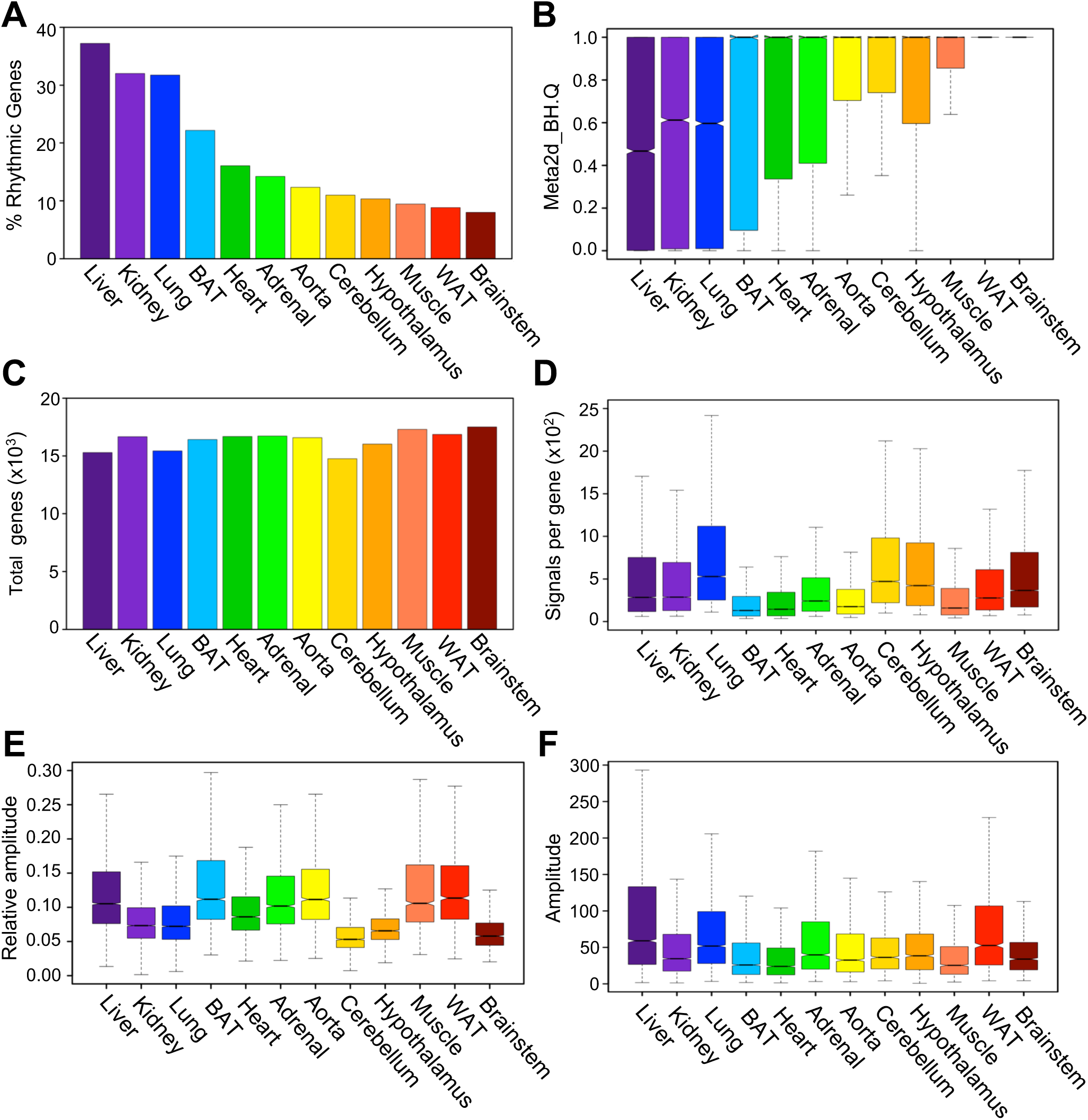
Characteristics of the mouse circadian transcriptome (microarray) in various mouse tissues. (A) Percentage of cycling genes in each tissue from highest (left) to lowest (right) % cycling. Rhythmicity of a gene was defined as Benjamini-Hochberg q values < 0.05 by MetaCycle. (B) Distribution of Benjamini-Hochberg q values of all expressed genes in each tissue. (C) Numbers of genes expressed in each tissue. (D) Average microarray signals per gene for all the probesets. (E) Distribution of relative amplitude of cycling genes in each tissue calculated by MetaCycle. (F) Distribution of the amplitude of cycling genes in each tissue calculated by MetaCycle. (D-F) The central line represents the median, and each box represents the 25th and 75th percentiles, respectively. The notch represents the 95% confidence interval around the median. Numbers of expressed genes or rhythmic genes in each tissue can be found in the Supplementary Data Sheet 1. Each color corresponds to a tissue; liver (purple), kidney (light purple), lung (blue), brown adipose (BAT) (light blue), heart (green), adrenal (light green), aorta (yellow), cerebellum (gold), hypothalamus (orange), muscle (coral), white adipose (WAT) (red), and brainstem (dark red).

Differences in the percentage of cycling genes in each tissue were not due to differences in the “strength” of the transcriptome. First, the total number of transcripts detected in each tissue was comparable, irrespective of the percentage of rhythmic transcripts (Fig. 1C). Second, the median microarray signals per transcript varied among tissues, and did not correlate with the percentage of rhythmic transcripts (Pearson R^2^=0.115, p=0.368; Spearman *rho*=-0.042, p=0.904) (Fig. 1D). When we focused only on the transcripts that were rhythmically expressed (Benjamini-Hochberg q<0.05), the median of the relative amplitude (i.e., the ratio between amplitude and baseline expression level) (Fig. 1E) or the amplitude itself (Fig. 1F) of cycling gene expression did not correlate with the percentage of cycling genes in each tissue (relative amplitude: Pearson r^2^=0.060, p=0.189; Spearman *rho*=0.070, p=0.834, amplitude: Pearson r^2^=0.439, p=0.154; Spearman *rho*=0.175, p=0.588), indicating that the amplitude of gene expression is comparable in each tissue if they are rhythmic.

We also performed the same set of analyses using the RNA-seq data ^10^, which surveyed the same set of tissues but with a lower time resolution (Microarray: 2 hrs, RNA-seq: 6 hrs) ^10^. Even though the order of the tissues was slightly different than the microarray datasets (Fig. S1A), which was likely due to the differences in time resolution and the threshold to eliminate low-expressed transcripts (see Materials and Methods), the results were essentially the same: distribution of Benjamini-Hochberg q-values from the rhythmicity analysis was wider in tissues with a high percentage of rhythmic transcripts (Fig. S1B), the strength of transcriptome was comparable among tissues (Fig. S1C, D), and the median of the relative amplitude or the amplitude of cycling gene expression was comparable in each tissue (Fig. S1E, F).

### Characterization of cycling gene expression in various mouse tissues

Because we did not observe any characteristics that had a correlation with the percentage of cycling genes at a genome-wide scale, we shifted our focus on single gene level analyses. We first analyzed a total of 15 core clock genes (*Arntl, Clock, Npas2, Per1-3, Cry1-2 Rora-c, Nr1d1- 2, Dbp, Nfil3*), and found that most of these genes were expressed ubiquitously across all tissues, except for *Rorb*, whose expression was restricted to brain and brown adipose tissue (Supplemental Data File 1a and 1b). A majority of these genes were also rhythmically expressed, except for the *Rors*: *Rora* was rhythmic in four tissues (liver, lung, heart, and muscle) but not in the other eight tissues (kidney, BAT, adrenal, aorta, cerebellum, hypothalamus, WAT, and brainstem), *Rorb* was arrhythmic in all 4 tissues that it is expressed in (BAT, hypothalamus, cerebellum, and brainstem), and *Rorc* was rhythmic in most of the peripheral tissues but not in the hypothalamus, muscle, or brainstem (Supplemental Data File 1), which was consistent with previous reports ^7,9,29,30^. *Clock* and *Cry1* were also arrhythmic in hypothalamus, making the hypothalamus the tissue with the lowest number of rhythmic core clock genes, even though hypothalamus ranked 9th out of 15 tissues in the percentage of cycling transcripts (Fig. 1A). Similar results were obtained from the RNA-seq data, although the number of rhythmic core clock genes were lower, most likely due to the lower time resolution of the RNA-seq data compared to the microarray data (Supplemental Data File 2).

As was previously reported, the phases of core clock gene expression were confined to a relatively narrow window ^12,30-33^, except for a few genes such as *Cry2, Rorc*, and *Nfil3* (Fig. 2A). On the other hand, the relative amplitude of core clock gene expression was more variable between tissues (Fig. 2B), and nine genes, *Dbp, Npas2, Nr1d1, Arntl, Per3, Per2, Rorc, Cry1, and Cry2* had their relative amplitude positively correlated with the percentages of cycling transcripts in either Pearson and Spearman correlation analyses (Table 1). Additional ten clock-controlled genes were expressed and rhythmic in all tissues, for which we calculated the correlation between their relative amplitude and the percentage of cycling transcripts. Among those, the relative amplitude of three genes (*P4ha1, Tsc22d3*, and *Lonrf3*), or another set of three genes (*Tsc22d3, Lonrf3*, and *Usp2*) correlated significantly with the percentage of cycling transcripts in Spearman or Pearson analyses, respectively (Table 1). Notably, the strongest correlation was observed for *Rorc* (Pearson r^2^=0.852, p=0.004; Spearman *rho*=0.917, p=0.001). It is unclear, however, whether the higher amplitude of core clock gene expression leads to a higher percentage of rhythmic transcripts in each tissue or *vice versa*. We also analyzed the data from RNA-seq and performed the same analyses. However, the lower number of rhythmic core clock genes detected in the RNA-seq dataset significantly compromised our ability to calculate the correlations between the percentage of rhythmic transcripts in each tissue and the phase and amplitude of core clock gene expression.

**Table 1.** Correlations between the percentage of rhythmic transcripts and the relative amplitude of core clock genes in each tissue

|  | Pearson (linear) |  | Spearman (non-linear) |  |
| --- | --- | --- | --- | --- |
|  | R<sup>2</sup> | P-value | *Rho* | P-value |
| *Arntl* | 0.5775568 | 0.04923* | 0.8741259 | 0.0003089* |
| *Clock* | 0.3962219 | 0.2277 | 0.3181818 | 0.3414 |
| *Npas2* | 0.6054532 | 0.04839* | 0.7181818 | 0.0168* |
| *Per1* | 0.4538226 | 0.1384 | 0.4685315 | 0.2175 |
| *Per2* | 0.6617942 | 0.01907* | 0.7972028 | 0.0031161* |
| *Per3* | 0.5899605 | 0.04347* | 0.5944056 | 0.04575* |
| *Cry1* | 0.7952259 | 0.003433* | 0.7090909 | 0.01873* |
| *Cry2* | 0.5342694 | 0.07355 | 0.6853147 | 0.01731* |
| *Nr1d1* | 0.6248975 | 0.02981* | 0.5944056 | 0.04575* |
| *Nr1d2* | 0.5542365 | 0.06149 | 0.4405594 | 0.1542 |
| *Rora* | 0.3681567 | 0.5421 | 0.2 | 0.7833 |
| *Rorc* | 0.8522986 | 0.003518* | 0.9166667 | 0.001312* |
| *Dbp* | 0.6452423 | 0.02346* | 0.6853147 | 0.01731* |
| *Nfil3* | 0.4639099 | 0.1506 | 0.3272727 | 0.327 |
| *P4ha1* | 0.5738972 | 0.05103 | 0.6923077 | 0.01588* |
| *Tsc22d3* | 0.677488 | 0.01549* | 0.6293706 | 0.03239* |
| *Lonrf3* | 0.74423 | 0.005506* | 0.7972028 | 0.003161* |
| *Usp2* | 0.5768827 | 0.04956* | 0.4685315 | 0.1275 |

\*Asterisks denote p<0.05
\*\* *Rorb* was excluded from our correlation analyses due to its low expression in all tissues except brain

**Figure 2.**
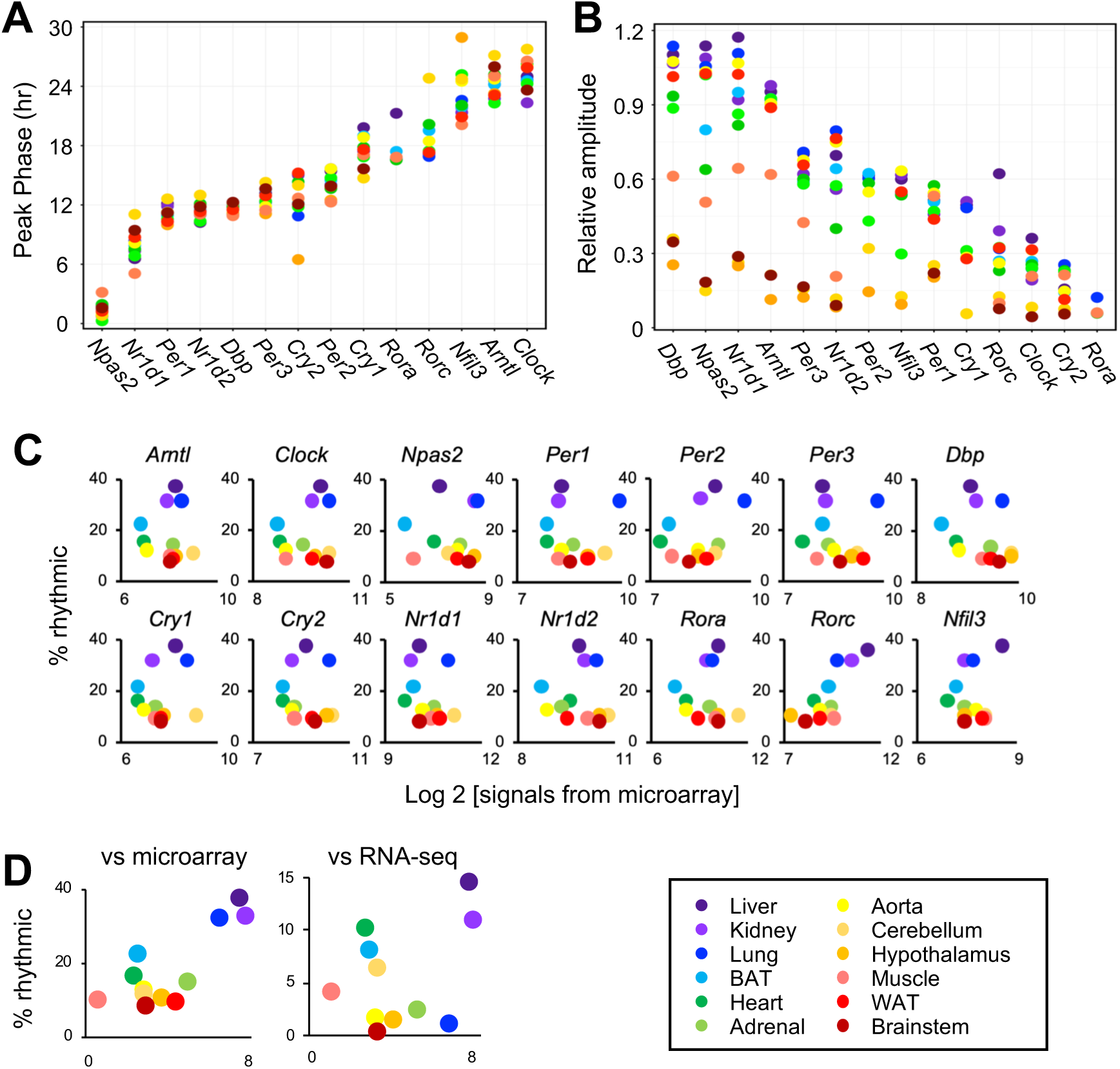
A positive correlation between the percentage of cycling transcripts and the mean level of *Rorc*. (A) The peak phase of core clock gene expression in CT (Circadian Time) determined by MetaCycle. (B) The relative amplitudes of core clock gene expression determined by MetaCycle. (C) Correlation between % rhythmic transcripts and the mean levels of each core clock gene determined by MetaCycle. (D) Correlation between % rhythmic transcripts and the mean levels of *Per2AS* in each tissue. Each color corresponds to a tissue; liver (purple), kidney (light purple), lung (blue), brown adipose (light blue), heart (green), adrenal (light green), aorta (yellow), cerebellum (gold), hypothalamus (orange), muscle (coral), white adipose (red), brainstem (dark red). Core clock gene expression that did not fulfill Benjamini-Hochberg q < 0.05 criteria for rhythmicity was not included.

We also investigated the correlation between the mean levels of core clock gene expression across all time points and the percentage of cycling genes in each tissue, as non-cycling genes could contribute to the differences in the percentage of cycling genes. Interestingly, we again found that there was a positive correlation between the percentage of cycling transcripts and the mean level of *Rorc*, but not with any other core clock genes (Fig. 2C, Table 2). This correlation was also observed in the RNA-seq dataset (Fig. S2, Table 2). We also tested *Per2AS*, a newly identified non-coding RNA ^34-36^, because the expression of *Per2AS* is rhythmic and antiphasic to *Per2* in liver, adrenal gland, lung, and kidney ^10^, and it was suspected that *Per2AS* is involved in regulating the circadian system ^37^. Indeed, we found a linear correlation between the mean levels of *Per2AS* and the percentages of rhythmic transcripts in both microarray and RNA-seq datasets (Fig. 2D).

**Table 2.** Correlations between the percentage of rhythmic genes and the mean expression level of core clock genes in microarray and RNA-seq datasets

|  | Pearson (linear) |  |  |  | Spearman (non-linear) |  |  |  |
| --- | --- | --- | --- | --- | --- | --- | --- | --- |
|  | Microarray |  | RNA-seq |  | Microarray |  | RNA-seq |  |
|  | R<sup>2</sup> | P-value | R<sup>2</sup> | P-value | *rho* | P-value | *rho* | P-value |
| *Arntl* | 0.00025806 | 0.9993649 | -0.2477924 | 0.46255259 | -0.0629371 | 0.8459309 | -0.3545455 | 0.28469274 |
| *Clock* | 0.20709174 | 0.5184007 | -0.3077513 | 0.35722663 | -0.020979 | 0.9484022 | -0.3818182 | 0.24655958 |
| *Npas2* | 0.13268917 | 0.6810054 | NA | NA | 0.03496503 | 0.9140933 | NA | NA |
| *Per1* | 0.07460386 | 0.8177602 | -0.4232656 | 0.19458479 | -0.2797203 | 0.3785687 | -0.4454545 | 0.1697326 |
| *Per2* | 0.4444646 | 0.1477156 | -0.3037159 | 0.36389063 | 0.25174825 | 0.4299188 | -0.1 | 0.769875 |
| *Per3* | -0.004058 | 0.9900137 | -0.5366997 | 0.08871493 | -0.3006993 | 0.3422595 | -0.6181818 | 0.04264557 |
| *Cry1* | 0.17502327 | 0.5863932 | -0.1887732 | 0.57828653 | -0.1258741 | 0.6966831 | -0.4272727 | 0.18994372 |
| *Cry2* | -0.0725671 | 0.8226649 | -0.469578 | 0.14504423 | -0.3636364 | 0.245265 | -0.6454545 | 0.0319628 |
| *Nr1d1* | -0.1931836 | 0.5474633 | -0.4214839 | 0.19667007 | -0.4055944 | 0.1908359 | -0.5818182 | 0.0604199 |
| *Nr1d2* | -0.1093403 | 0.7351637 | -0.3684187 | 0.26490747 | -0.3006993 | 0.3422595 | -0.5090909 | 0.10973723 |
| *Rora* | -0.154493 | 0.6316449 | -0.0850269 | 0.8037002 | -0.2097902 | 0.5128409 | -0.4 | 0.22286835 |
| *Rorc* | 0.81551562 | 0.0012236* | 0.80572369 | 0.0027537* | 0.65734266 | 0.0201855* | 0.8 | 0.0031104* |
| *Dbp* | -0.3043041 | 0.3362164 | -0.5439689 | 0.08366 | -0.4615385 | 0.1309481 | -0.4454545 | 0.1697326 |
| *Nfil3* | 0.2775392 | 0.3824536 | 0.18797349 | 0.57992718 | -0.1188811 | 0.7128842 | 0.4 | 0.22286835 |
| *Per2AS* | 0.8546421 | 0.0003983* | 0.6079273 | 0.04723* | 0.4545455 | 0.1404 | 0.07272727 | 0.8388 |

\*Asterisks denote p<0.05
\*\* *Rorb* was excluded from our correlation analyses due to its low expression in all tissues except brain

To test how robust the correlation of *Rorc* is, we extended the analysis to the genome-wide scale. We found that among 12,024 genes expressed in all 12 tissues from microarray datasets, the mean level of 1,131 and 400 genes was correlated significantly with the percentage of rhythmic genes in each tissue from the Pearson (linear) and the Spearman (non-linear) correlation test, respectively (Supplemental Data File 3). Similarly, among 8,269 genes expressed in all 11 tissues from the RNA-seq datasets, we found the mean level of 925 and 1,664 genes correlated significantly with the percentage of rhythmic genes from the Pearson and Spearman correlation tests, respectively (Supplemental Data File 3). Of those, 135 (Pearson) or 77 (Spearman) genes were found correlated in both microarray and RNA-seq datasets (Supplemental Data File 3), and we therefore considered those as more robustly correlated. Gene ontology (GO) analyses were then performed to assess whether a specific process contributes to the high percentage of rhythmic transcripts. No pathways were detected as statistically significant (FDR < 0.05) among those that correlated robustly in the Spearman analysis. Whereas numerous metabolic processes were enriched among those that correlated robustly in the Pearson analysis (Supplemental Data File 4). We also calculated Fisher Z-scores from each test to evaluate the relative strength of *Rorc* correlation, compared to other genes. *Rorc* was ranked 5th (Pearson) or 14^th^ (Spearman) when we used average Z-scores from both microarray and RNA-seq datasets. These data suggest that the correlation between the level of *Rorc* and the amplitude of the mouse circadian transcriptome is one of the strongest.

### The effect of RORC as a transcriptional activator in regulating the circadian transcriptome

Because *Rorc* directly activates the transcription of RORE-containing genes ^29,31,38^, we hypothesized that, if *Rorc* was directly driving rhythmic gene expression leading to a high number of cycling transcripts, then the number of rhythmic genes with RORE motifs in their promoter would be higher in tissues with a higher number of rhythmic genes. To test this, we first retrieved the promoter sequence of rhythmic genes in each tissue using the RNA-seq dataset (−1000 bp to +100 bp with respect to TSS), and then determined the number of DNA motifs that can be recognized by RORC *in silico*. We surveyed the recognition sequences of not only RORC, but also RORA, RORB, NR1D1, NR1D2, ARNTL, CLOCK, NPAS2, NFIL3, and DBP, because these proteins are all considered important to drive rhythmic gene expression ^39,40^. We found that the RORC binding motif was found in approximately 8% of the rhythmically expressed genes, and this was consistent in all 11 tissues we examined (Fig. 3). The binding sites for NPAS2, ARNTL, NR1D1, and NR1D2 were the most highly represented (∼10-14 %), and NFIL3 and DBP were the least represented (∼3-6 %) (Fig. 3). These data indicate that the effect of RORC is most likely indirect and that the level of *Rorc* does not directly contribute to driving rhythmic gene expression. Instead, the level of *Rorc* is probably important to increasing the amplitude of circadian transcriptional output in the circadian system as a whole, and this ultimately results in increasing the number of cycling genes.

**Figure 3.**
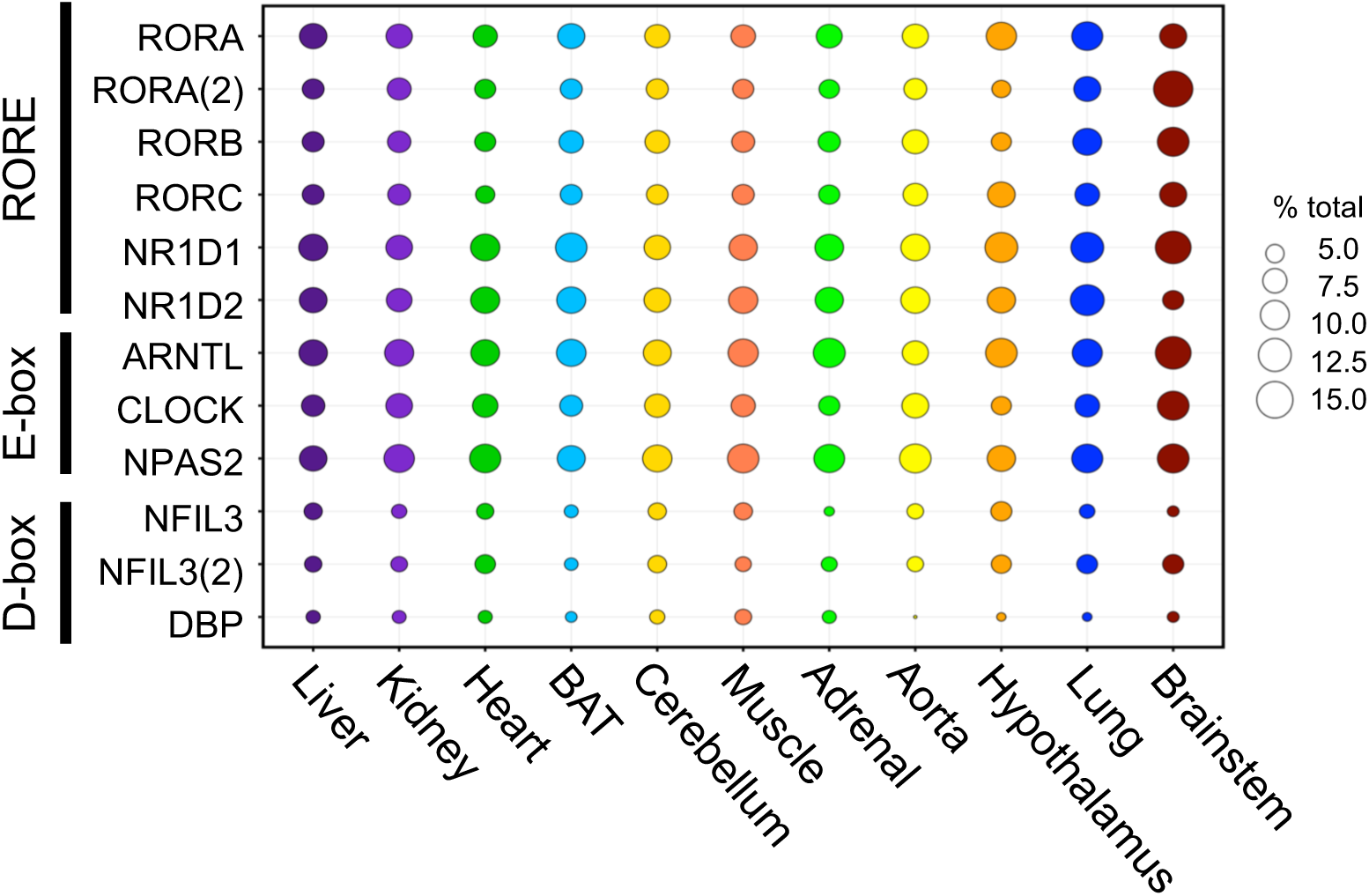
The number of RORC-binding motifs does not correlate with the percentage of rhythmic genes in each tissue. Weighted scatterplot representing the percentage of rhythmic genes containing binding motifs of circadian transcription factors listed on the left. The size of each circle represents the % and each color corresponds to a tissue.

### Potential mechanism of RORC in regulating the amplitude of the circadian transcriptome

To further gain more mechanistic insights into how *Rorc* contributes to the increase in the number of cycling mRNAs without driving mRNA expression, we next tested the correlation between the mean levels of *Rorc* and other core clock genes. Not surprisingly, we found a linear correlation between the mean levels of *Rorc* and *Per2AS* in both the microarray and RNA-seq datasets (Table 3). The mean level of *Rorc* also linearly correlated with *Nfil3* (Microarray) or *Cry2* (RNA-seq) (Table 3); however, the biological significance of these correlations is unclear, as the correlations were not consistent between microarray and RNA-seq datasets. We did not detect any correlation between *Rorc* and *Rorc*-target genes such as *Arntl, Cry1, Nfil3*, and *Nr1d1* (Table 3), whose promoter regions contain RORE motifs and the amplitude of their rhythmic mRNA expression was dampened in most of the *Rorc*^*-/-*^ tissues ^40-43^. We did not find any correlations between the level of *Per2AS* and other core clock genes either, except for *Per2* and *Npas2* (Table 4). The significance of these correlations is also unclear, because they were found only in microarray but not in RNA-seq datasets.

**Table 3.** Correlations between the mean level of *Rorc* and the mean level of other clock genes in each tissue

|  | Pearson (linear) |  |  |  | Spearman (non-linear) |  |  |  |
| --- | --- | --- | --- | --- | --- | --- | --- | --- |
|  | Microarray |  | RNA-seq |  | Microarray |  | RNA-seq |  |
|  | R<sup>2</sup> | P-value | R<sup>2</sup> | P-value | *Rho* | P-value | *Rho* | P-value |
| *Arntl* | 0.1424678 | 0.6587 | -0.1930034 | 0.5696 | 0.2517483 | 0.4301 | -0.3 | 0.3711 |
| *Clock* | 0.2017612 | 0.5295 | -0.5247839 | 0.09743 | 0.2307692 | 0.4709 | -0.5636364 | 0.07594 |
| *Npas2* | -0.1087769 | 0.7341 | -0.1029352 | 0.8262 | 0 | 1 | -0.1071429 | 0.8397 |
| *Per1* | -0.1271442 | 0.6938 | -0.3823291 | 0.2459 | 0.02097902 | 0.9562 | -0.5727273 | 0.0706 |
| *Per2* | 0.2802252 | 0.3777 | 0.03352763 | 0.922 | 0.5244755 | 0.08388 | 0 | 1 |
| *Per3* | -1.93E-01 | 0.5478 | -0.5847199 | 0.05884 | -0.1328671 | 0.6834 | -0.7818182 | 0.007012 |
| *Cry1* | 0.2039933 | 0.5248 | -0.3352892 | 0.3135 | 0.2237762 | 0.4849 | -0.5909091 | 0.06073 |
| *Cry2* | -0.1406772 | 0.6628 | -0.7071451 | 0.01495* | -0.020979 | 0.9562 | -0.8 | 0.005202 |
| *Nr1d1* | -0.1865557 | 0.5615 | -0.2836711 | 0.3979 | -0.013986 | 0.9737 | -0.5636364 | 0.07594 |
| *Nr1d2* | -0.075446 | 0.8157 | -0.6073742 | 0.4749 | -0.0489511 | 0.8863 | -0.6818182 | 0.02548 |
| *Rora* | 0.08273569 | 0.7982 | -0.2710108 | 0.4202 | 0.0979021 | 0.7663 | -0.4545455 | 0.1634 |
| *Dbp* | -0.2537079 | 0.4262 | -0.2642677 | 0.4323 | -0.1608392 | 0.6194 | -0.4272727 | 0.1926 |
| *Nfil3* | 0.6783749 | 0.01531* | 0.4541654 | 0.1605 | 0.4195804 | 0.1766 | 0.4727273 | 0.1456 |
| *Per2AS* | 0.858603 | 0.0003493* | 0.6431022 | 0.0328* | 0.5034965 | 0.09875 | 0.1454545 | 0.6734 |

\*Asterisks denote p<0.05
\*\* *Rorb* was excluded from our correlation analyses due to its low expression in all tissues except brain

**Table 4.** Correlations between the mean *Per2AS* TPM and the mean level of other clock genes in each tissue

|  | Pearson (linear) |  |  |  | Spearman (non-linear) |  |  |  |
| --- | --- | --- | --- | --- | --- | --- | --- | --- |
|  | Microarray |  | RNA-seq |  | Microarray |  | RNA-seq |  |
|  | R<sup>2</sup> | P-value | R<sup>2</sup> | P-value | *Rho* | P-value | *Rho* | P-value |
| *Arntl* | 0.1072332 | 0.7401 | -0.1923029 | 0.5711 | 0.4405594 | 0.1542 | 0.4727273 | 0.1456 |
| *Clock* | 0.3049452 | 0.3351 | -0.102975 | 0.7632 | 0.5804196 | 0.05209 | 0.5272727 | 0.1001 |
| *Npas2* | 0.308457 | 0.3292 | -0.0209776 | 0.9644 | 0.6783217 | 0.01883* | 0.5357143 | 0.2357 |
| *Per1* | -0.0469417 | 0.8848 | -0.3423037 | 0.3028 | 0.4125874 | 0.1845 | 0.0090909 | 0.9892 |
| *Per2* | 0.3363529 | 0.2851 | 0.03514764 | 0.9183 | 0.7202797 | 0.01102* | 0.5636364 | 0.07594 |
| *Per3* | -6.68E-02 | 0.8367 | -0.2394287 | 0.4783 | 0.4265734 | 0.1689 | -0.0454546 | 0.9029 |
| *Cry1* | 0.1160046 | 0.7196 | -0.1523629 | 0.6547 | 0.4125874 | 0.1845 | 0.2090909 | 0.5391 |
| *Cry2* | -0.0904382 | 0.7798 | -0.360859 | 0.2756 | 0.2727273 | 0.3912 | 0.1090909 | 0.7549 |
| *Nr1d1* | -0.2317755 | 0.4685 | -0.4086626 | 0.2121 | 0.1258741 | 0.6997 | -0.0545455 | 0.8815 |
| *Nr1d2* | 0.03833355 | 0.9058 | -0.2832654 | 0.3986 | 0.1188811 | 0.7162 | -0.0181818 | 0.9676 |
| *Rora* | -0.027873 | 0.9315 | -0.1883284 | 0.5792 | 0.2447552 | 0.4435 | 0.1363636 | 0.6935 |
| *Dbp* | -0.1331303 | 0.68 | -0.2910061 | 0.3853 | 0.2937063 | 0.3543 | 0.08181818 | 0.8177 |
| *Nfil3* | 0.357542 | 0.2539 | -0.1930106 | 0.5696 | 0.1468531 | 0.6511 | -0.3909091 | 0.2365 |

\*Asterisks denote p<0.05
\*\* *Rorb* was excluded from our correlation analyses due to its low expression in all tissues except brain

## Discussion

Among the three key parameters for cycling systems (period, phase, and amplitude), the regulatory mechanisms of period and phase have been relatively well-characterized, whereas that of amplitude have remained much more enigmatic. Forward genetics or screening approaches using pharmacological or genetic perturbation have not been successful, as the variance of amplitude is much higher than that of period, compromising the statistical ability to distinguish true positives from false positives ^44-48^. Furthermore, it is currently unclear whether it is a single gene, a combination of genes, one of the feedback loops, and/or topology of the network that is important for amplitude. To make things even more complicated, amplitude can be measured by various outputs, such as gene expression, firing patterns in neurons, body temperature, and locomotor activity, each of which can be under the regulation of both cell-autonomous (intracellular or local) and systemic (or extracellular) rhythms. It also remains unclear whether all the amplitude of various rhythms is regulated by the same mechanism.

In this study, we focused on the percentage of cycling genes in various mouse tissues and explored the possible mechanisms of amplitude regulation of circadian transcriptomic output. The circadian transcriptome can be influenced both by cell-autonomous and systemic cues in each tissue. However, the circadian gene expression is a common feature of the circadian clock system in various tissues and allows us to directly compare the difference in amplitude between tissues without relying on their respective physiology.

We found that 18 genes (eight core clock genes and ten clock-controlled genes) are rhythmically expressed in all tissues that we examined (Table 1, Supplemental Data File 3). This is consistent with previous observations that the rhythmicity of each gene is often tissue-specific, and while only a handful of genes are cycling in all or most tissues, others are rhythmic only in certain tissues ^7-11^. In addition, the relative amplitude of 13 genes (nine core clock genes and four clock-controlled genes) were correlated with the percentage of cycling genes, while the mean level of 197 genes correlated with the percentage of cycling genes. These genes are not necessarily expressed rhythmically, albeit about a half (100/197) are, and vast majority of these genes were involved in the metabolic processes (Supplemental Data File 3, 4). Given that the energy cost for cycling genes are higher than non-cycling genes ^49^, it is reasonable that metabolism related genes are highly expressed in tissues that have higher number of cycling transcripts. Nevertheless, it is unlikely that clock-controlled genes directly regulate or contribute to determine the number of cycling genes, as these processes are under the circadian control (i.e., output pathway). Rather, it is tempting to postulate that the role of *Rorc* and/or *Per2AS* in the core-clock circuit gives it a more promising function in potentially regulating the amplitude of the circadian transcriptome, at least in the tissues where *Rorc* is expressed, as the correlation of *Rorc* in both relative amplitude and the average level was one of the strongest. It is unclear from our study whether these are simply a correlation or causation and this needs to be studied in the future. Furthermore, it would also be of great interest to test whether the RORC protein levels, not mRNAs, also correlate with the amplitude of circadian transcriptome.

We also found that the effect of *Rorc* is most likely independent of its activity as a transcriptional activator (Fig. 3). Rather, *Rorc* appears to be one of the components that regulate the amplitude of the circadian system, particularly the part that is important for the circadian transcriptome (if different parts regulate different amplitude). The positive loop (*Clock-Arntl- Rev-Ror*) was originally considered to confer additional robustness to the system and, therefore, stabilizes the system. However, it is not required for circadian rhythm generation ^42,50^. Recent studies have also highlighted the role of the positive loop as the central axis of amplitude regulation ^51,52^. Our study is consistent with these findings and suggest that the positive loop, particularly the level of *Rorc*, is important in setting the amplitude of the circadian transcriptome. In addition, our study also suggested that *Per2AS* is involved in the positive loop, because the level of *Per2AS* positively correlated with the level of *Rorc* as well as the percentage of cycling genes in each tissue, even though it was originally assumed to only interact with *Per2* ^37^. Interestingly, the mathematical model predicted the functional interaction between *Ror* and *Per2AS*, as *Per2AS* would restore stable circadian rhythms when they are disrupted by the overproduction of *Ror* or *Rev-erb* mRNAs ^37^. It is possible that *Rorc* and *Per2AS* function synergistically in the circadian clock system.

It still remains unclear why *Rorc*, but not *Rora* and *Rorb*, correlates with the amplitude of the circadian transcriptome, as all the ROR proteins share significant sequence similarities ^38,53^. Unfortunately, the physiological relevance of each ROR paralogue has never been clarified in the mammalian circadian system. One notable difference among *Ror* paralogues, however, is their unique expression patterns (Supplementary Data Files 1a, 1b, 2). It is possible that the systemic cues, which are, in theory, the same to all the tissues have a tissue-specific impact in regulating the level of *Rorc*. Understanding the difference in the regulatory mechanisms of *Ror* gene expression would provide insight into how their tissue-specific expression pattern is achieved and how *Rorc* specifically impacts the amplitude of the circadian transcriptomic output.

Overall, our study highlighted the potential role of *Rorc* in regulating the amplitude of the circadian transcriptome. Follow-up experimental studies would further complement our observations from the rich transcriptomic datasets that are publicly available and delineate the mechanisms of circadian amplitude regulation.

## Acknowledgement

The authors thank Drs. Frank Aylward (Department of Biological Sciences, Virginia Tech) and Natalia Toporikova (Department of Biology, Washington and Lee University) for technical assistance and Dr. Janet Webster (Fralin Life Sciences Institute, Virginia Tech), as well as the members of the Kojima laboratory, for critical reading of the manuscript. This work was supported by R01GM126223 from the NIH (to S.K.).

## Author Contributions Statement

ESL performed data analysis, SK designed and conceived the study, and both ESL and SK generated figures, wrote, and edited the manuscript.

## Data Availability Statement

The datasets generated during and/or analyzed during the current study are available in the NCBI GEO repository, from series GSE54650.

