## Supplementary Information for "Genome-wide correlation analysis reveals *Rorc* as potential amplitude regulator of circadian transcriptome output"

Supplemental Figure 1. Littleton and Kojima

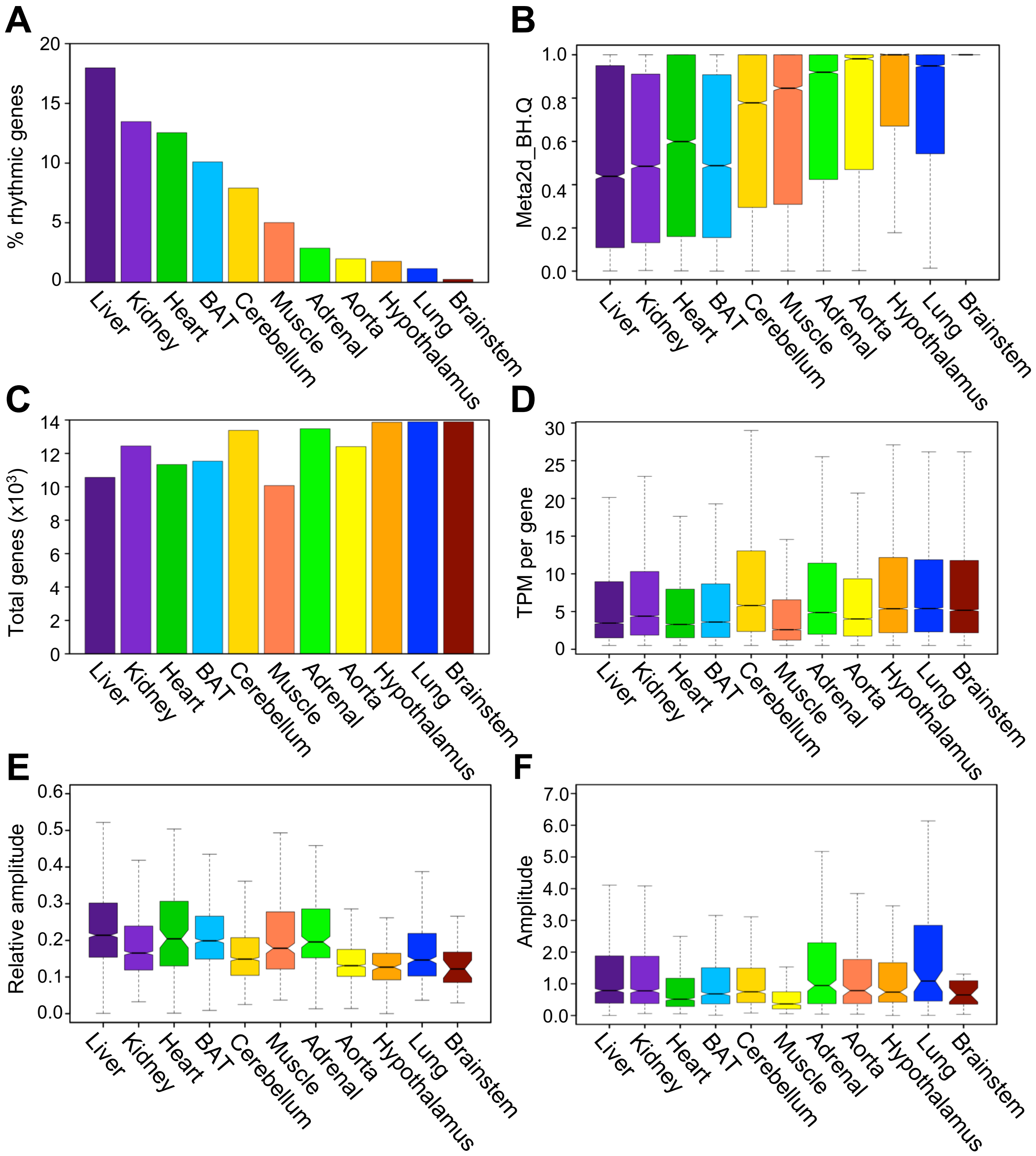

Supplemental Figure 2. Littleton and Kojima

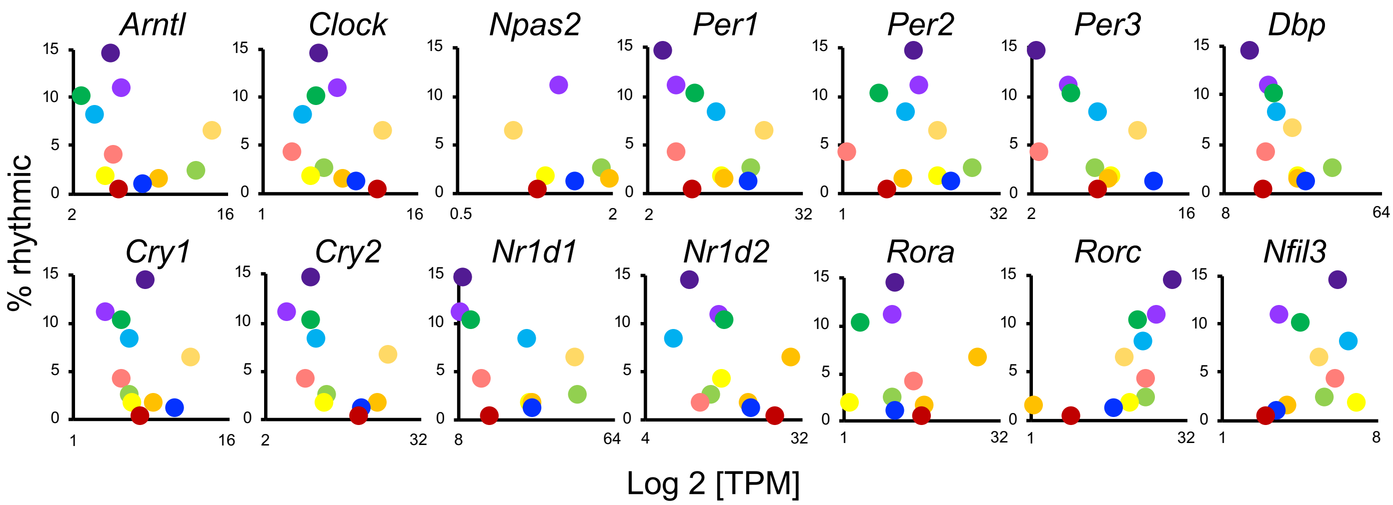

**Supplemental Figure Legends**

**Figure S1. Characteristics of mouse circadian transcriptome (RNA-seq) in various mouse tissues.** (A) Percentage of cycling genes in each tissue from highest (left) to lowest (right) % cycling. Rhythmicity of a gene was defined as Benjamini-Hochberg q values < 0.05 by MetaCycle. (B) Distribution of Benjamini-Hochberg q values of all expressed genes in each tissue. (C) Numbers of genes expressed in each tissue. (D) Average microarray signals per gene for all probesets. (E) Distribution of relative amplitude of cycling genes in each tissue calculated by MetaCycle. (F) Distribution of the amplitude of cycling genes in each tissue calculated by MetaCycle. (D-F) The central line represents the median, and each box represents the 25th and 75th percentiles, respectively. The notch represents a 95% confidence interval around the median. Numbers of expressed genes or rhythmic genes in each tissue can be found in the Supplementary Data Sheet 2. Each color corresponds to a tissue; liver (purple), kidney (light purple), lung (blue), brown adipose (BAT) (light blue), heart (green), adrenal (light green), aorta (yellow), cerebellum (gold), hypothalamus (orange), muscle (coral), and brainstem (dark red).

**Figure S2. Correlation between the percentage of cycling genes and the expression patterns of core clock genes in each tissue in RNA-seq data.** Colors of each tissue correspond to Fig. 1; liver (purple), kidney (light purple), lung (blue), brown adipose (light blue), heart (green), adrenal (light green), aorta (yellow), cerebellum (gold), hypothalamus (orange), muscle (coral), white adipose (red), brainstem (dark red).
